# Topologically frustrated dynamics in an uncharged Tetra-PEG gel

**DOI:** 10.1101/2022.02.23.481633

**Authors:** Di Jia, Yui Tsuji, Mitsuhiro Shibayama, Murugappan Muthukumar

**Affiliations:** Department of Polymer Science and Engineering, University of Massachusetts, Amherst, Amherst, Massachusetts 01003, United States; Beijing National Laboratory for Molecular Sciences, State Key Laboratory of Polymer Physics and Chemistry, Institute of Chemistry Chinese Academy of Sciences, Beijing 100190, China; University of Chinese Academy of Sciences, Beijing 100049, China; Institute for Solid State Physics, The University of Tokyo, Kashiwanoha, Kashiwa 277-8581, Japan; Neutron Science and Technology Center, Comprehensive Research Organization for Science and Society, Tokai, Ibaraki 319-1106, Japan

## Abstract

Based on our previous discovery of non-diffusive topologically frustrated dynamics in a charged system where electrostatic interactions between the charged guest and the charged host gel play a role (***Nature Communications***, 2018, 9, 2248; ***Physical Review Letters***, 2021, 126, 057802), we have investigated the onset of this effect in an uncharged gel matrix. Using sodium (polystyrene sulfonate) as the guest macromolecule and the ideal tetra-PEG gel, we find the emergence of the non-diffusive topologically frustrated dynamical state with a hierarchy of segmental dynamics represented by a stretched exponential of exponent β around 1/3. Our results demonstrate the universal behavior of the topologically frustrated dynamical state.

## 1. Introduction

Movement of very long polymer chains in restricted environments is ubiquitous in separation science and associated technologies, as well as in crowded biological situations [1-23]. While Brownian motion and diffusion of these chains at finite temperature is the axiom of their motion at ambient conditions, a recent experimental discovery [11,12] based on the dynamics of DNA and synthetic polyelectrolytes, trapped inside a hydrogel at room temperature containing essentially water, has revealed that this axiom is wildly deviated at intermediate confinements. The guest polymers are plunged into an extremely long-lived metastable dynamical state, called the topologically frustrates dynamical state. The reason behind this phenomenon has been conjectured to arise from elicitation of numerous entropic traps into which a very long polymer chain is distributed. The hierarchical non-diffusive dynamics observed in the new dynamical state is also conjectured to arise from either polydispersity in the mesh size of the hydrogel or conformational fluctuations of the portions of the chain in various meshes. In view of this, we address in this paper the role of distribution of mesh size of the gel on the onset of the topologically frustrated dynamical state. To reach this goal, we have monitored the dynamics of guest poly(styrene sulfonate) molecules trapped inside ideal networks, with narrow mesh size, made from tetra-polyethylene glycol. We report that the topologically frustrated dynamical state still emerges in these ideal gels, showing that the origin of the new phenomenon is due to conformational fluctuations of the various sectors of the guest macromolecule trapped inside the meshes of the host gel.

## 2. Experimental section

### 2.1. Materials

Tetra-amine terminated poly(ethylene glycol) (Tetra-PEG-NH_2_) with the molecular weight M_w_=40kDa and tetra-NHS-glutarate-terminated poly(ethylene glycol) (Tetra-PEG-NHS) with the molecular weight M_w_=40kDa were purchased from NOF Co. (Tokyo, Japan). Here NHS represents N-hydroxysuccinimide. Sodium polystyrene sulfonate (NaPSS) with the molecular weight M_w_=2270kDa was purchased from Scientific Polymers. Sodium phosphate, disodium phosphate, and sodium chloride were purchased from Sigma-Aldrich. Hydrophilic Polyvinylidene Fluoride (PVDF) filters with the pore size 450 nm were purchased from Millex Company. Deionized water was obtained from a Milli-Q water purification system (Millipore, Bedford, MA, U.S.A.). The resistivity of deionized water used was 18.2 MΩ cm.

### 2.2. Gel Synthesis and Sample Preparation

Two types of 4-armed poly(ethylene glycol) (PEG) macromonomers with equal arm length and with different reactive terminal groups were used to synthesize Tetra-PEG gels. Tetra-PEG gels were synthesized by cross-end-coupling of N-succinimidyl (NHS) terminated tetra-PEG macromonomers and amine-terminated tetra-PEG macromonomers following the previous method^1-4^. Tetra-PEG-NH_2_ and Tetra-PEG-NHS were dissolved in phosphate buffer to adjust the pH so that pH=7.4 for Tetra-PEG-NH_2_ solution and pH=5.8 for Tetra-PEG-NHS solution. The ionic strength of the buffer solution was tuned by adding NaCl. Phosphate buffer with ionic strength 100 mM (pH=6.8) was used as the solvent. Equimolar quantities of Tetra-PEG-NH_2_ and Tetra-PEG-NHS were mixed with certain amount of concentrated PSS solutions to reach the targeted concentrations for each species. The polymer concentration of the Tetra-PEG gel was 60 mg/mL and the NaPSS concentration was 0, 5, 20, and 40 mg/mL, respectively. Since light scattering measurement is extremely sensitive to dust, the light scattering tubes were first washed with pure water and acetone separately for several times. After they were dried in the oven overnight, aluminum foil was used to wrap up the tubes and then these tubes were further cleaned by distilled acetone through an acetone fountain setup^5,6^. Then the pre-gel solutions were slowly filtered into a light scattering tube through a 450 nm PVDF filter to remove the dust. All the sample preparation work was conducted in a super-clean bench to avoid the dust. At least 24 hours were allowed for the completion of the reaction before the light scattering measurements were performed.

### 2.3. Static and dynamic light scattering

SLS and DLS measurements were performed on a commercial spectrometer equipped with a multi-τ digital time correlator (ALV-5000/E) with Argon laser light source (output power = 400 mW) of wavelength 514.5 nm. DLS measures the intensity-intensity time correlation function g_2_(t) by means of a multi-channel digital correlator and related to the normalized electric field correlation function g_1_(t) through the Siegert relation^7^. Measurement was conducted at the scattering angle 30°, 40°, 50°, 60°, and 70°, respectively. The relaxation time was obtained by averaging over three different spatial locations within the samples.

### 2.4 Data fitting method

CONTIN method and multiple exponential fitting method were used to analyze the characteristic relaxation rate Γ at each angle^7-10^. Based on D=Γ/q^2^, diffusion coefficient D was calculated. Here q is the scattering wavevector q = (4πn/λ)sin(θ/2) with the scattering angle θ, wavelength of the incident monochromatic light λ, and refractive index of the medium n. For the correlation functions of NaPSS in the gel matrix, multiple modes were first confirmed by CONTIN method. In the presence of multiple modes and in order to facilitate comparison to a theory, the normalized electric field correlation function g_1_(t) was fitted by a sum of one or two exponential decays and a stretched exponential function^11-13^. g_1_(t) was fitted using ORIGIN software by minimization of errors between the fitted prediction and the data. Iterations were performed until the best fitting curve was obtained within the tolerance limit. The residuals, which are obtained from the difference between the original data and the fitting curve, are randomly distributed about the mean of zero and do not have systematic fluctuations about their mean.

## 3. Results and discussion

### 3.1 Characterization of the gel matrix

The gel matrix used here was an uncharged Tetra-PEG gel with monodispersed gel mesh size, because the Tetra-PEG gel was made from two types of 4-armed PEG macromonomers with equal arm length crosslinked through their end groups, as shown in Figure 1(e). Static and dynamic light scattering measurements were conducted to characterize the gel matrix. The normalized correlation function g_1_(t) and the corresponding relaxation time distribution function obtained from CONTIN fit at the scattering angle 30o as a typical example is shown in Figure1(a) and 1(b). Typically for gels, the correlation function obtained from DLS shows a dominant decay with several other relaxations due to inherent inhomogeneities from clustering of cross-links and other impurities^13-16^. These inhomogeneities show up as a static electric field, which interferes directly with the scattered electric field from the gel mode. However, here there is only one mode, indicating the gel is very homogenous and clean. And the gel mode is diffusion-like representing the gel elasticity, and the corresponding elastic diffusion coefficient D_gel_ is (4.37±0.15)E-7 cm^2^/s (Figure1c). From static light scattering measurement, the gel mesh size is obtained by using Ornstein-Zernike equation^7^:

**Figure 1.**
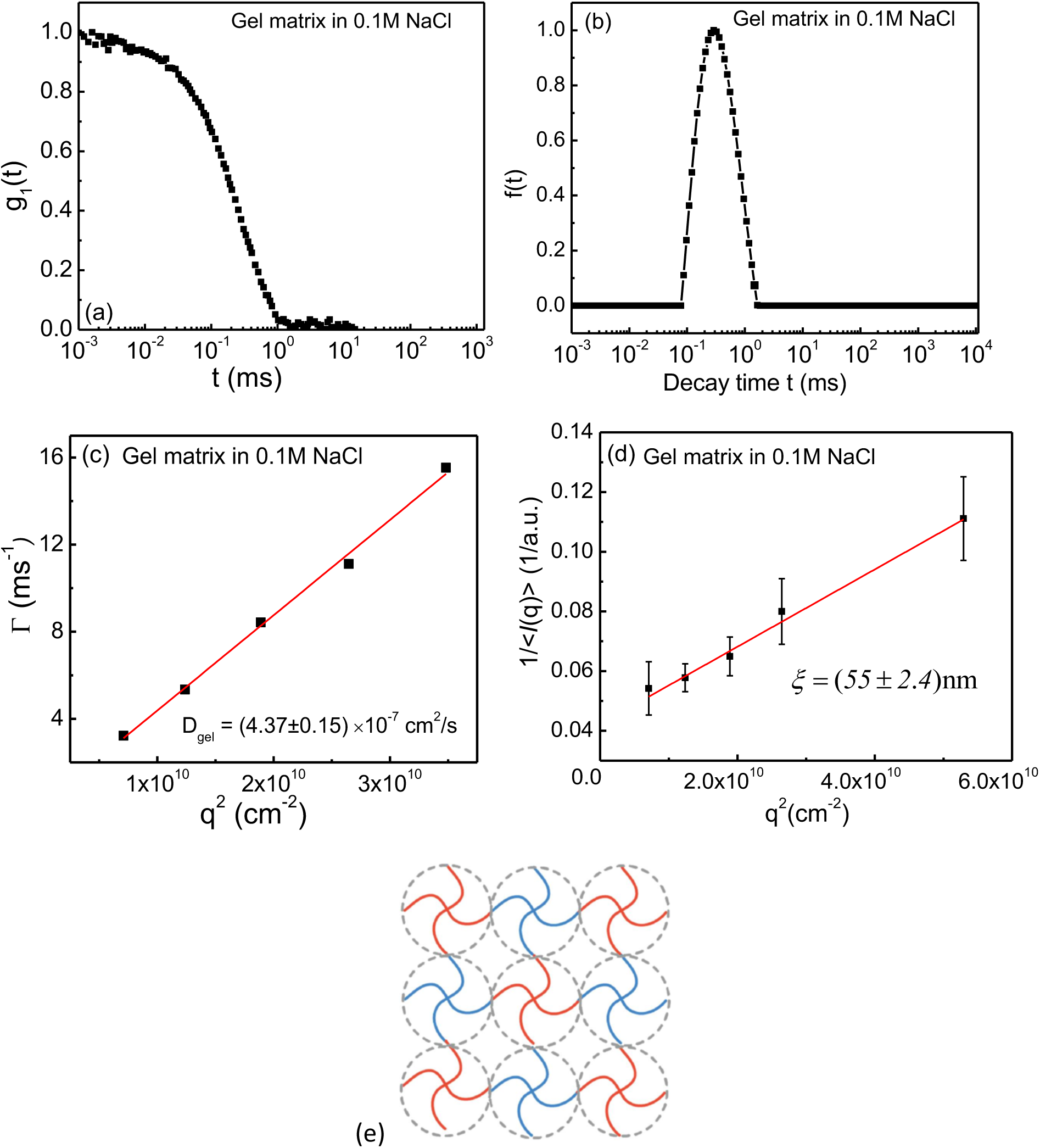
Dynamic and static light scattering results for the Tetra-PEG gel matrix with 0.1M NaCl. (a) Normalized field-correlation function g_1_(t) at scattering angle 30^°;^ measured by dynamic light scattering. (b) Corresponding relaxation time distribution function obtained from CONTIN fit at scattering angle 30^°;^. (c) q^2^ dependence of the relaxation rate Γ for the Tetra-PEG gel matrix. (d) Ornstein-Zernike plot of the inverse averaged scattering intensity 1/<I(q)> versus q^2^ measured by static light scattering of the Tetra-PEG gel matrix, yielding the averaged mesh size ξ= (55±2.4) nm. (e) Schematic cartoon of the Tetra-PEG gel with monodispersed gel mesh size synthesized by crosslinked the end groups of two types of 4-arm macromonomers with equal arm length.

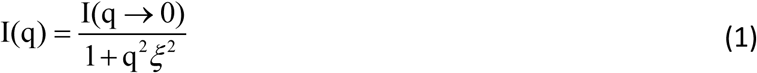

Where I(q) is the static light scattering intensity at the scattering wave vector q, ξ is the correlation length of the gel and here it is used to estimate the gel mesh size. In the plot of 1/I(q) vs q^2^, we get ξ= (55±2.4)nm from the square root of the slope/intercept ratio, as shown in Figure 1(d). It is to be noted that the mesh size determined from Ornstein-Zernicke analysis is different from that obtained from DLS, since the latter does not measure the mesh size, but only elastic moduli.

### 3.2 Dynamics of guest chains in the Tetra-PEG gel matrix at high concentration

The guest chain used here is NaPSS with M_w_=2270kDa in 0.1M NaCl solution. The detailed characterization of NaPSS with M_w_=2270KDa in 0.1M NaCl solution is in our previous studies^10,17^. For NaPSS with M_w_=2270KDa in 0.1M NaCl solution, the SLS results show that its radius of gyration is Rg=96nm. The DLS results show that there are three diffusive modes representing coupled motion between couterions, co-ions, polyions, and dipole-dipole interaction induced aggregation mode. The key point here is when NaPSS chains are in solutions, all the modes are diffusive. However, it is entirely different when the NaPSS guest chains were imbedded inside the gel matrix.

The best fit of the correlation function g_1_(t) for 40g/L NaPSS in the gel matrix can be expressed as (figure 2a):

**Figure 2.**
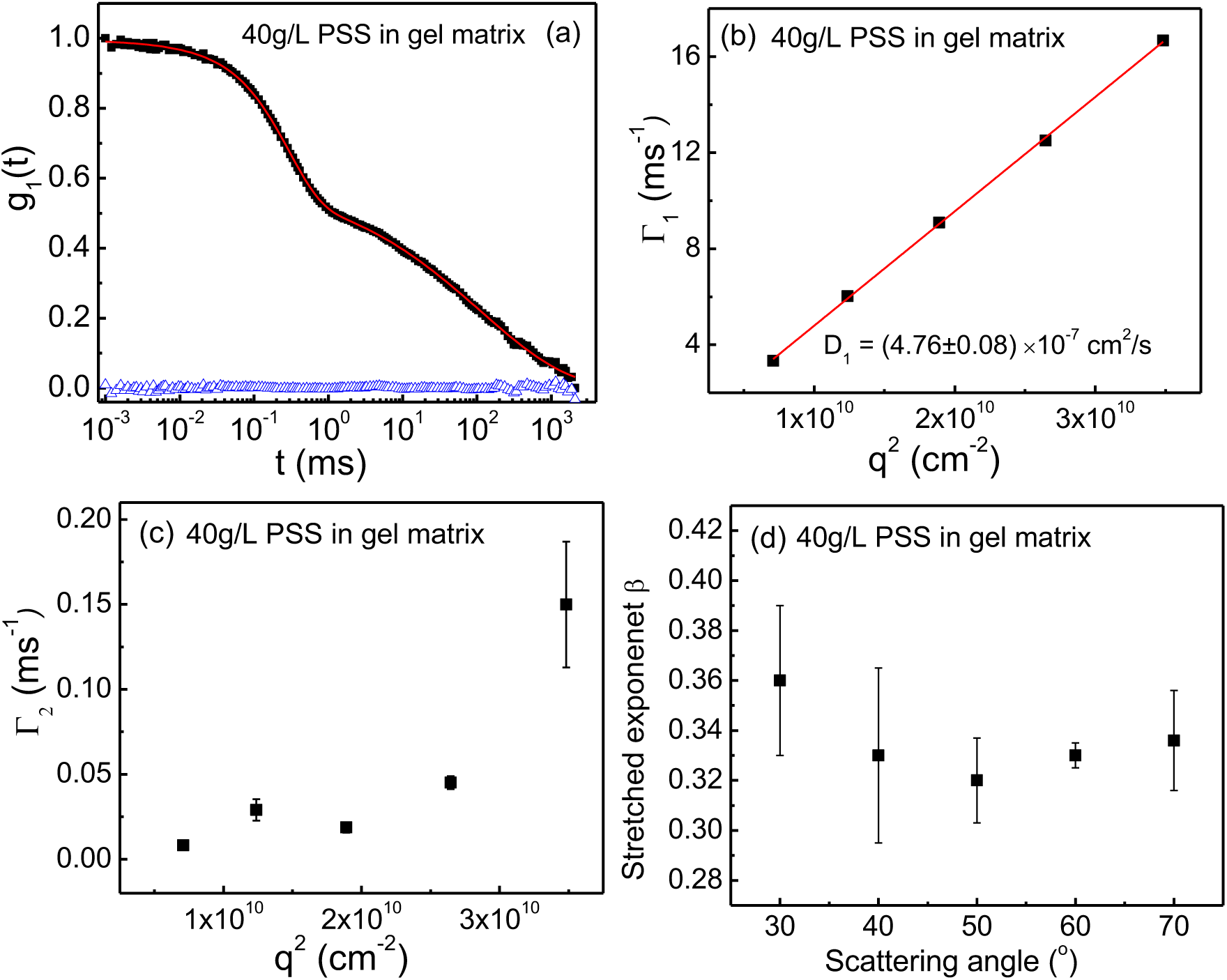
Dynamics of 40g/L NaPSS in the Tetra-PEG gel matrix with 0.1M NaCl. (a) Normalized field-correlation function g_1_(t) at scattering angle 30°. The blue triangles are the residuals between the original data (black) and the fitting curves (red). (b), (c) Fitting results of the relaxation rate Γ vs. q^2^ at all angles for the first and second mode, respectively. (d) Stretched exponent β as a plot of different scattering angles for 40g/L NaPSS in the Tetra-PEG gel matrix with 0.1M NaCl.

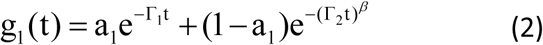

where Γ_1_=0.29ms^-1^ and Γ_2_=112.82 ms^-1^ are the decay rates for the two dynamical modes, a_1_= 0.4 is the weight of the first mode, and the value of the exponent β= 0.36 is a measure of non-diffusive hierarchically dynamical cooperativity of PSS chains in the gel matrix. Analyzing g_1_(t) data at multiple scattering angles reveals that the first mode (exponential decay) is diffusive, which is demonstrated in Figure 2(b) where Γ_1_∼q^2^ with the diffusion coefficient D_1_= (4.76±0.08) E-7cm^2^/s. Since D_1_ is very close to the diffusion coefficient of the gel matrix alone (Figure1c), so we assign the first mode as the gel mode, which indicates the gel elasticity (D_1_=D_gel_). Then the dynamics of all the guest chains come to the second term in Eq (2), which is a non-diffusive stretched exponential decay, because Γ_2_ is not proportional to q^2^ (Figure 2c). And the value of β, which indicates the hierarchy of the cooperative local segmental chain dynamics, is independent of scattering angles (Figure 2d), and the averaged value of β over all scattering angles is β= (0.33±0.04). The non-diffusive stretched exponential decay of the guest chain dynamics under confinement is ascribed to the non-diffusive topological frustrated dynamics, in which the center of mass of the whole chain cannot diffuse, but within one chain all the segments can do their local segmental dynamics in a hierarchical way. And the β values at all scattering angles are within a very narrow range, which is very different from the traditional temperature-dependent glassy and aging systems, where β is in the full range (0<β<1)^18^. It is also remarkable that all the three diffusive modes in solutions are replaced by a non-diffusive stretched exponential mode when the guest chains are inside the gel matrix.

The discovery here is different from our previous studies, since the previous studies are all for the charged system with a negative charged poly (acrylamide-co-sodium acrylate) gel matrix and either negatively or positively charged polyelectrolytes as the guest chains (***Nature Communications***, 2018 9, 2248; ***Phys. Rev. Lett***. 2021, 126, 057802). While in this work, we have used an uncharged Tetra-PEG neutral gel matrix and NaPSS as the guest chains so that electrostatic interactions between the guest chains and the host gel matrix can be avoided. But still we are able to observe the non-diffusive topologically frustrated dynamics in an uncharged gel matrix, demonstrating such a phenomenon is universal and is ubiquitous for both charged systems and uncharged systems.

Besides, the mesh size distribution of the Tetra-PEG gel matrix is highly monodispersed due to its special synthetic method, while the mesh size distribution of the charged poly(acrylamide-co-sodium acrylate) gel is rather polydispersed because it is synthesized by the traditional free radical polymerization^19,20^. And previously we think that the polydispersity of the mesh size in the gel matrix is the possible reason for the emergence of the non-diffusive topologically frustrated dynamics. But here we have observed the non-diffusive topologically frustrated dynamics in a highly monodispersed uncharged gel matrix, demonstrating the emergence of the non-diffusive topologically frustrated dynamics is not due the polydispersity of the mesh size in the gel matrix.

### 3.3 Evidence of weak complexation between NaPSS and the Tetra-PEG gel matrix

When the concentration of the guest chain in the Tetra-PEG gel is low (5g/L), there are strong fluctuations for the scattered light intensity (Figure 3a) and such strong scattered light intensity fluctuations can easily destroy the scattered light intensity-intensity correlation function g_2_(t) so that we cannot obtain the valid correlation function g_2_(t)^21-23^. As shown in Figure 3(a), for 5 g/L NaPSS in the gel matrix, the averaged scattered intensity Is is 30 (a.u.), while the maximum Is is 120 (a.u.), which is 4 times larger than the averaged value. Such strong fluctuations of Is can immediately destroy the scattered light intensity-intensity correlation function g_2_(t), so that we cannot get the valid g_2_(t) from the measurement. On the other hand, for high concentration of NaPSS in the gel, such as 40 g/L and 20g/L, fluctuations in Is becomes smaller by increasing the guest molecules, allowing to obtain a valid scattered light intensity-intensity correlation function g_2_(t), as shown in Figure 3(b) and 3(c).

**Figure 3.**
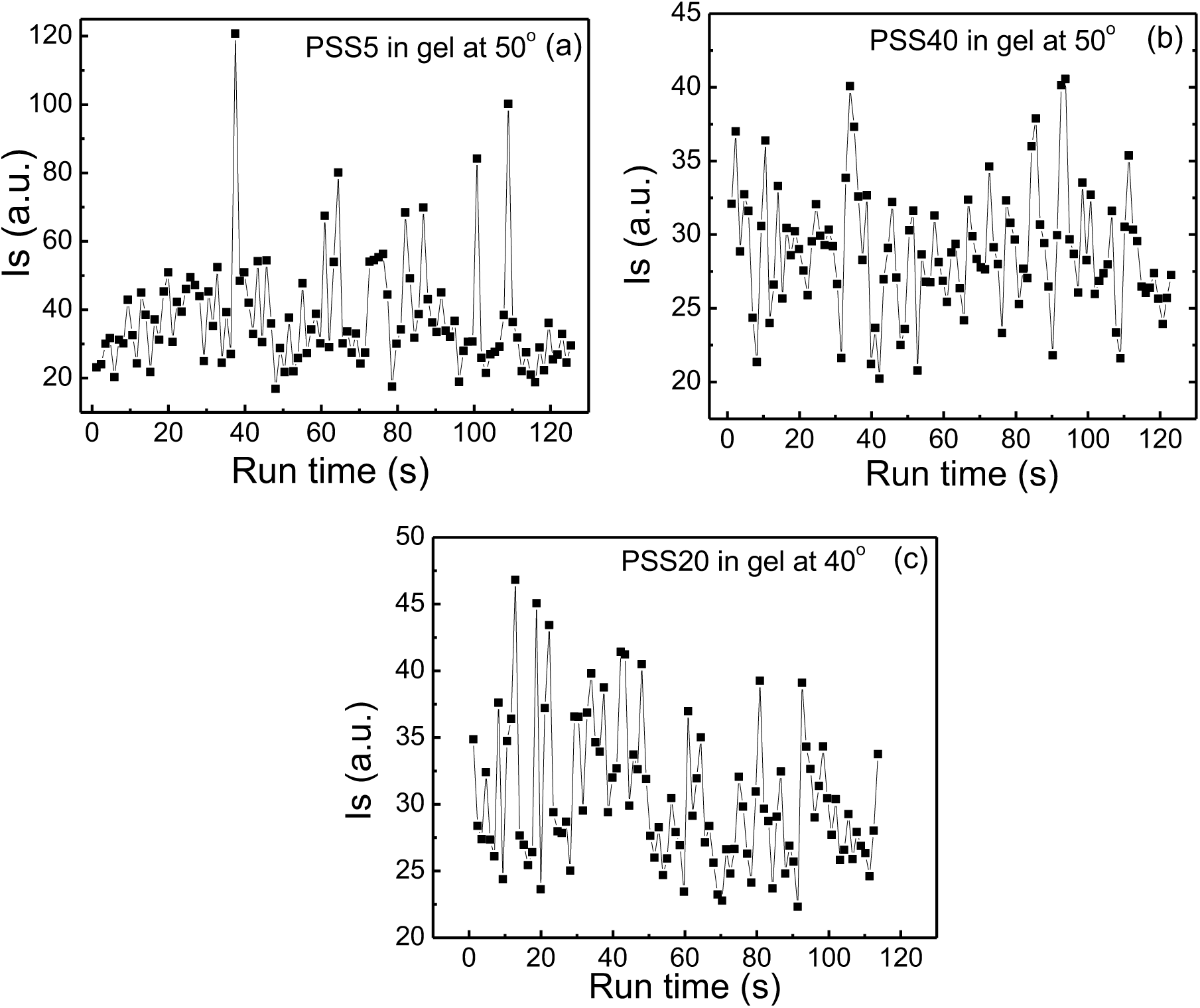
Scattered intensity Is as a function of running time during the measurements for NaPSS of 5 g/L (a), 20 g/L (b), 40 g/L(c) in the gel matrix, respectively.

Therefore, we came up with the hypothesis that there is weak complexation between NaPSS chains and the Tetra-PEG gel matrix and such weak complexation can induce strong scattered light intensity fluctuations. When NaPSS concentration is low, all the NaPSS can weakly complex with Tetra-PEG gel because there are enough complexation sites on the gel. However, when NaPSS concentration is high, only part of NaPSS can complex and fully saturated the complexation sites on the Tetra-PEG gel, then the extra free NaPSS chains can move around with no complexation, and the Is signal from the extra free NaPSS chains will dominate over the Is signal from the weakly complexed NaPSS chains. Therefore, we can successfully observe the topologically frustrated dynamics for the extra uncomplexed guest chains.

In order to verify our hypothesis, we have designed an experiment to test the diffusion coefficient D of each species and their mixtures in solutions as follows. We have measured the diffusion coefficients for 5g/L Tetra-PEG-NH_2_ (M_w_=40KDa) macromonomer solution, 5g/L Tetra-PEG-NHS macromonomer (M_w_=40KDa) solution, 10g/L NaPSS (M_w_=68KDa) solution, and their mixtures. The concentrations are chosen so that each species is below their overlap concentration C*. All the samples are with 0.1M NaCl. The results are shown in Table 1 and Figure 4. For 10g/L NaPSS (M_w_=68KDa) in 0.1M NaCl solution, there is only one mode with the diffusion coefficient D=(5.01±0.13)E-7 cm^2^/s. For two types of macromonomers, D=(3.88±0.06)E-7 cm^2^/s for 5g/L Tetra-PEG-NH_2_ in 0.1M NaCl solution and D=(3.87±0.05)E-7 cm^2^/s for 5g/L Tetra-PEG-NHS in 0.1M NaCl solution. When 10g/L NaPSS mixed with 5g/L Tetra-PEG-NHS solution, D=(4.05±0.02)E-7 cm^2^/s, which is between D of NaPSS alone and D of Tetra-PEG-NHS alone. However, when 10g/L NaPSS mixed with 5g/L Tetra-PEG-NH_2_ solution, D=(2.71±0.03)E-7 cm^2^/s, which is lower than both D of NaPSS alone and D of Tetra-PEG-NH_2_ alone, indicating there are bigger aggregates with lower diffusion coefficient show up due to weak complexation between NaPSS and Tetra-PEG-NH_2_. This is the evidence of the weak complexation between the NaPSS and the Tetra-PEG-NH_2_. When Tetra-PEG-NH_2_ macromonomers become the crosslinked gel, the complexation between the NaPSS and the gel made of Tetra-PEG-NH_2_ cannot form aggregates because now Tetra-PEG-NH_2_ species is static and frozen in the gel network and cannot freely move. Therefore, the weak complexation between NaPSS and the gel made of Tetra-PEG-NH_2_ can only lead to strong fluctuations of the scattered light intensity.

**Table 1.** Summary of diffusion coefficient D of NaPSS (M_w_=68KDa) solution, macromonomer solutions, and the mixture of NaPSS and macromonomer solutions. All the samples are with 0.1M NaCl.

| Sample name | Diffusion coefficient *D* |
| --- | --- |
| 10g/L NaPSS | (5.01±0.13)E-7 cm<sup>2</sup>/s |
| 5g/L Tetra-PEG-NH<sub>2</sub> | (3.88±0.06)E-7 cm<sup>2</sup>/s |
| 5g/L Tetra-PEG-NHS | (3.87±0.05)E-7 cm<sup>2</sup>/s |
| 10g/L NaPSS+5g/L Tetra-PEG-NH<sub>2</sub> mixture | (2.71±0.03)E-7 cm<sup>2</sup>/s |
| 10g/L NaPSS+5g/L Tetra-PEG-NHS mixture | (4.05±0.02)E-7 cm<sup>2</sup>/s |

**Figure 4.**
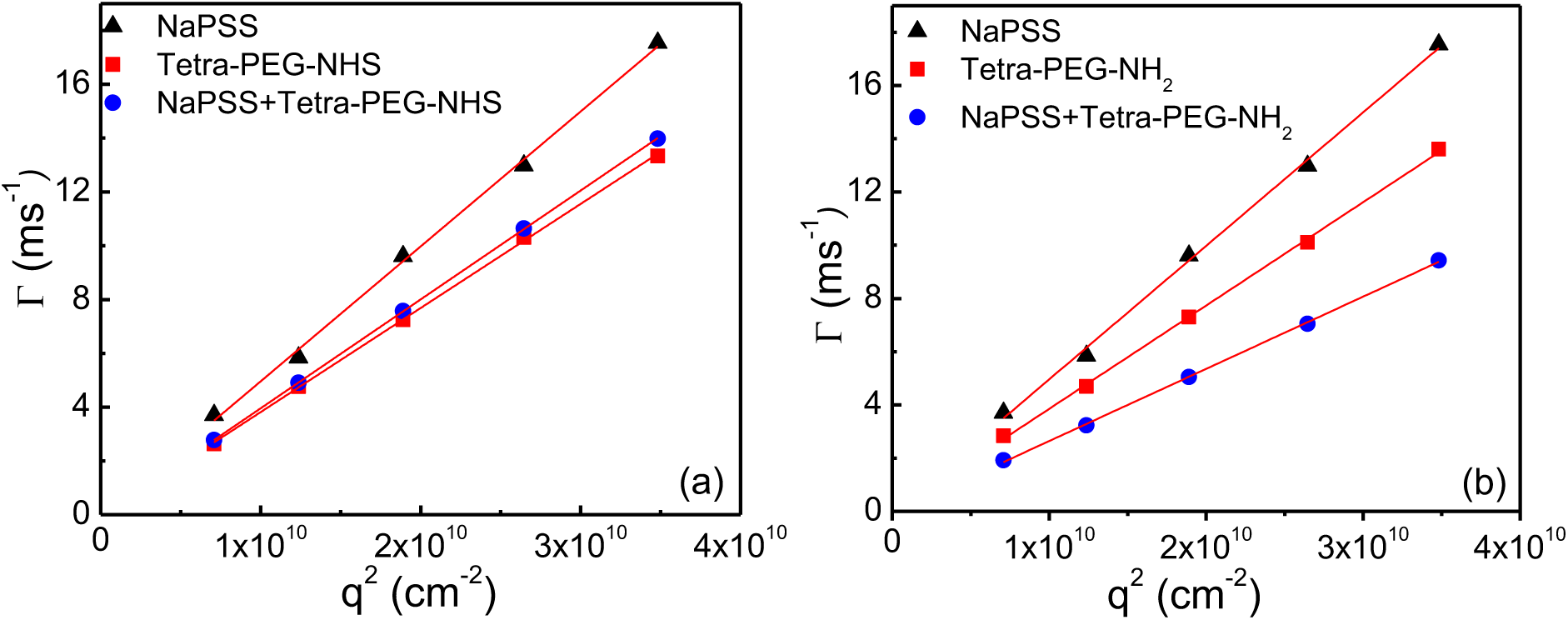
(a) q^2^ dependence of the relaxation rate Γ of 10mg/mL NaPSS (M_w_=68KDa) solution, 5mg/mL Tetra-PEG-NHS solution, and the mixture of 10mg/mL NaPSS and 5mg/mL Tetra-PEG-NHS solution. (b) q^2^ dependence of the relaxation rate Γ of 10mg/mL NaPSS (M_w_=68KDa) solution, 5mg/mL Tetra-PEG-NH_2_ solution, and the mixture of 10mg/mL NaPSS and 5mg/mL Tetra-PEG-NH_2_ solution. All the samples are with 0.1M NaCl.

### 3.4 Dynamics of the guest chains in the neutral gel matrix at intermediate concentration

Previous results (Figure 2) already show the NaPSS in the Tetra-PEG gel matrix at high concentration (40 g/L). Although we cannot obtain the valid scattered light intensity-intensity correlation function g_2_(t) measured by DLS at low NaPSS concentration (5 g/L) due to the strong fluctuations of the scattered light scattering. But we are still able to get the valid g_2_(t) at intermediate NaPSS concentration (20 g/L). The correlation function at the scattering angle 30o and 50^°^ are shown in figure 5(a) and (b) and the best fit of g_1_(t) can be accomplished by a sum of one exponential decay, one stretched exponential decay and another exponential decay with much larger decay time (ultra-slow mode), expressed as below:

**Figure 5.**
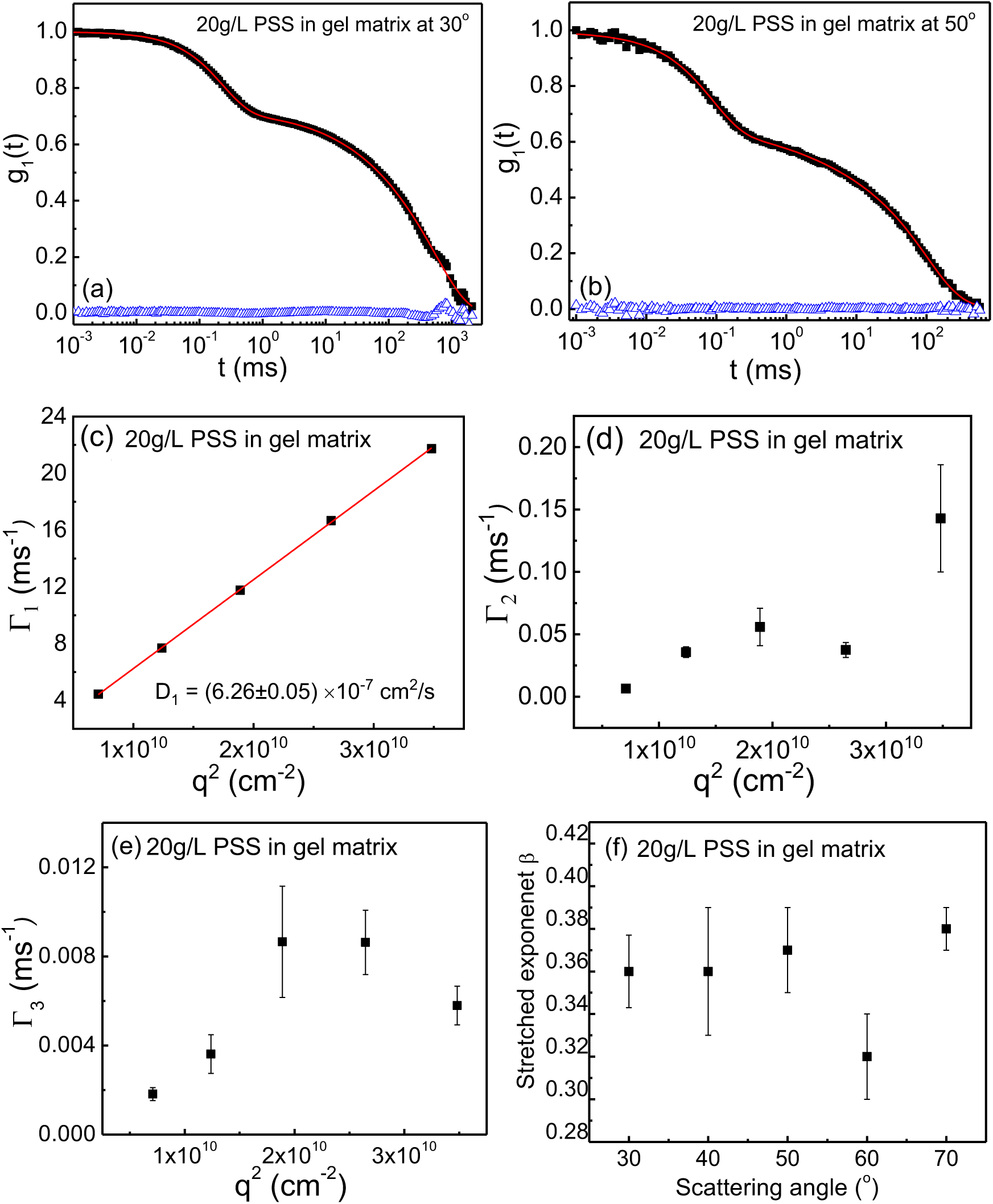
Dynamics of NaPSS chains at 20 g/L in the Tetra-PEG gel matrix with 0.1M NaCl. (a), (b) Normalized field-correlation function g_1_(t) at scattering angle 30° and 50° respectively. The blue triangles are the residuals between the original data (black) and the fitting curves (red). (c), (d), (e) Fitting results of the relaxation rate Γ_1_, Γ_2_, Γ_3_ vs. q^2^ at all angles for the first, second and third mode, respectively. (f) Stretched exponent β as a plot of different scattering angles for 20 g/L NaPSS in the Tetra-PEG gel matrix with 0.1M NaCl.

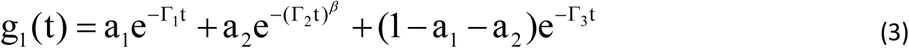

where for theg_1_(t) at scattering angle 30^°;^ (Figure5a), a_1_=0.25, Γ_1_=0.23ms^-1^, a_2_=0.4, Γ_2_=154.1ms^-1^, β=0.36, Γ_3_=550.3ms-1. And for the g_1_(t) at scattering angle 50^°^(Figure 5b), a_1_=0.31, Γ_1_=0.086ms^-1^, a_2_=0.38, Γ_2_=17.92ms^-1^, β=0.37, Γ_3_=115.5ms^-1^. The analysis of DLS data on the decay rates at different scattering angles shows that Γ_1_ is diffusive with the corresponding diffusion coefficient D_1_=(6.26±0.05)E-7 cm^2^/s, which is the gel mode (Figure 5c). The second mode is a non-diffusive stretched exponential decay since Γ_2_ is not proportional to q^2^ (Figure 5d). And the value of β over all the scattering angles is β=(0.36±0.03) ((Figure 5f). The third non-diffusive mode with a much slower decay rate Γ_3_ (ultra-slow mode) comes from the heterogeneity due to the weak complexation between NaPSS and Tetra-PEG gel matrix (Figure 5e). Since the value of Γ_3_ is much larger than Γ_2_ and Γ_1_, so luckily we can decouple the third ultra-slow mode due to heterogeneity from the second stretched exponential mode, which indicates the guest chain dynamics, although both the second and third modes are non-diffusive since both Γ_2_ and Γ_3_ are not proportional to q^2^.

## 4. Conclusions

As in our previous study using poly(acrylamide-co-acrylate) hydrogels as the host gel, where the mesh size of the gel is not uniform, we find the same phenomenon with similar non-diffusive hierarchical dynamics with the ideal tetra-PEG hydrogels, where the mesh size is uniform. This finding demonstrates that fluctuations in the mesh size of the matrix is not critical to elicit the topologically frustrated dynamical state. Furthermore, since we find the same value of β around 1/3 for both systems, we conclude that this new dynamical state is universal and ubiquitous to both charged and uncharged systems.

## Acknowledgements

MM acknowledges financial support from the National Science Foundation (DMR-2004493). DJ acknowledges financial support from Chinese Academy of Sciences.

